# On Neural Code – The Self-Information Processor disguised as neuronal variability?

**DOI:** 10.1101/132068

**Authors:** Joe Z. Tsien, Meng Li

**Affiliations:** Brain and Behavior Discovery Institute, Medical College of Georgia, Augusta University, Augusta, GA, USA; The Brain Decoding Center, Banna Biomedical Research Institute, Yunnan Academy of Science and Technology, Xishuangbanna, Yunnan, China

**Author notes:** Corresponding Author: Joe Z. Tsien.

## Abstract

One important goal of BRAIN projects is to crack the neural code — to understand how information is represented in patterns of electrical activity generated by ensembles of neurons. Yet the major stumbling block in the understanding of neural code is *neuronal variability* - neurons in the brain discharge their spikes with tremendous variability in both the *control* resting states and across trials within the same experiments. Such on-going spike variability imposes a great conceptual challenge to the classic rate code and/or synchrony-based temporal code. In practice, spike variability is typically removed via over-the-trial averaging methods such as peri-event spike histogram. In contrast to view neuronal variability as a noise problem, here we hypothesize that neuronal variability should be viewed as the *self-information processor*. Under this conceptual framework, neurons transmit their information by conforming to the basic logic of the statistical Self-Information Theory: spikes with higher-probability inter-spike-intervals (ISI) contain less information, whereas spikes with lower-probability ISIs convey more information, termed as *surprisal spikes*. In other words, real-time information is encoded not by changes in firing frequency per se, but rather by spike’s variability probability. When these surprisal spikes occur as positive surprisals or negative surprisals in a temporally coordinated manner across populations of cells, they generate cell-assembly neural code to convey discrete quanta of information in real-time. Importantly, such surprisal code can afford not only robust resilience to interference, but also biochemical coupling to energy metabolism, protein synthesis and gene expression at both synaptic sites and cell soma. We describe how this neural self-information theory might be used as a general decoding strategy to uncover the brain’s various cell assemblies in an unbiased manner.

---

With the emergence of various powerful technologies to aid brain research (1-5), neuroscientists are increasingly focusing on some of the biggest questions about the brain: What are the basic design principles underlying the brain’s wiring and computational logic (6-9)? How do groups of neurons convert raw sensory inputs into perception, memories, knowledge and actions (10-14)? How can real-time neural codes be constructed in the form of neural activity patterns that can be deciphered by both neurons and experimenters (15-25)?

While it is well established that neurons transmit information by using the number of spikes and/or the precise timing of these spikes — strategies often referred to as rate and/or temporal coding - in the actual brain, neurons fire spontaneously during both the “control” resting states and across different trials within the same experiment. Such on-going spike variability makes it difficult for rate-code (frequency code) to reliably decode stimulus identity in real-time (15, 26). Likewise, fluctuating spike discharge patterns also undermine synchrony-based rate code (27, 28). From structural perspective, it is not hard to see why such variability is inevitable. Neurons in the mammalian brain contain many thousands of synaptic connections, ranging from ∼30,000 synapses per pyramidal cells in the neocortex up to 200,000 synapses per purkinje cell in the cerebellum (29-33), with each synapse receiving ongoing inputs from active presynaptic cells (Figure 1A). Summation of these excitatory postsynaptic potentials (EPSP) triggers action potentials, or spikes, in the postsynaptic cell soma. With ongoing inputs coming from these tens of thousands of synapses, neurons ought to generate spike trains stochastically with enormous variability.

**Figure 1.**
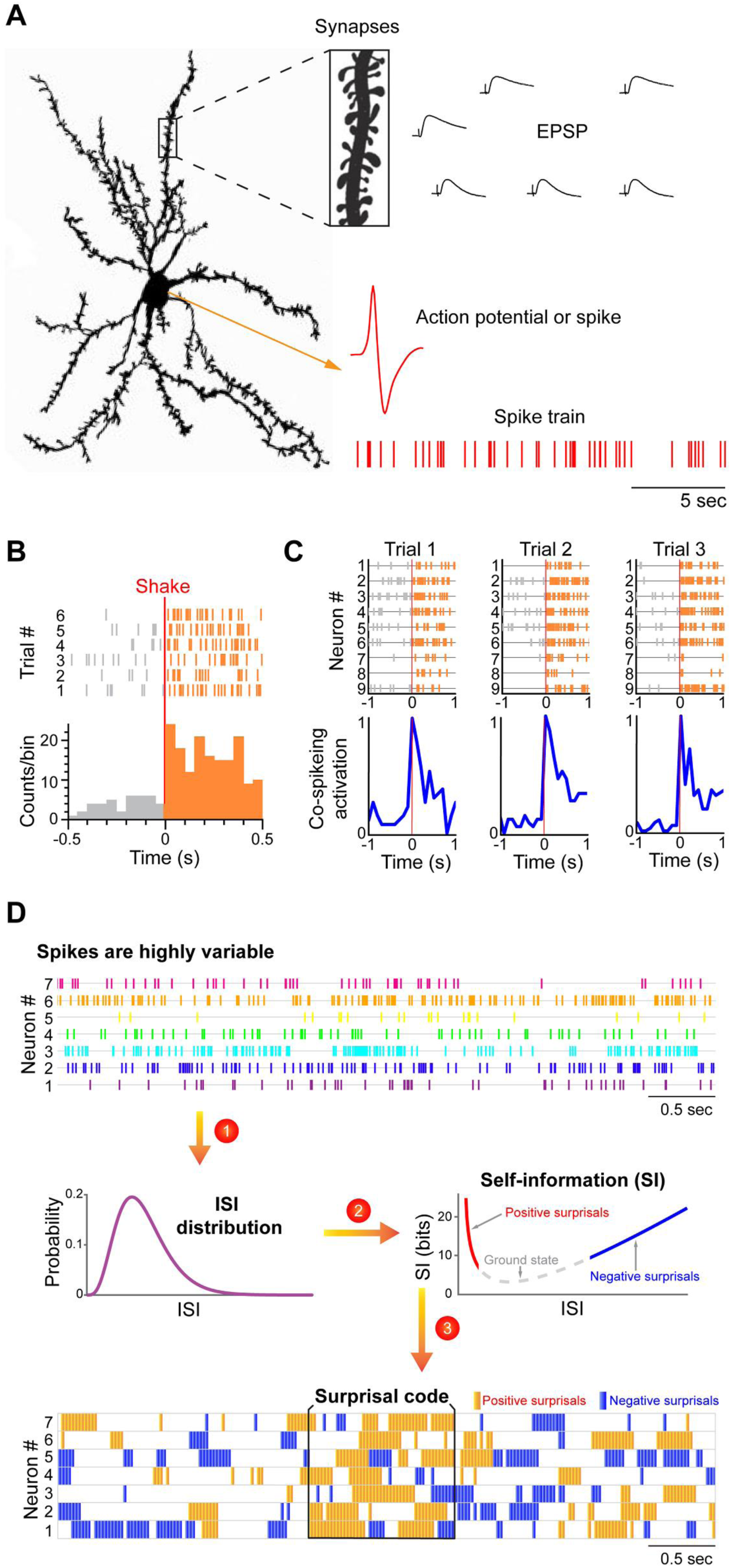
Neural Self-Information Theory provides a new way to understand how neural code is generated in the face of enormous spike variability. (A) A cortical neuron may contains ten thousands of synapses which can contribute to changes in excitatory post-synaptic potential (EPSP), leading to generation of action potential or spike. Stochastic nature of synaptic patterns lead to highly variable spike trains in both the resting “control” condition and stimulus-presentation experiments. (B) Response variability is typically removed or reduced by over-the-trial data averaging method such peri-event spike raster and histogram. A hippocampal neuron responded to earthquake (7 trials). (C) A group of the neurons with the similar response property, termed as neural clique, can be more effective to produce a more stable response. Nine anterior cingulate cortex cells responded to earthquake and formed the earthquake-specific neural clique. This neural clique showed robust response increase over three earthquake trials. (D) A general strategy to apply the neural self-information theory to uncover cell assemblies from spike train datasets. Surprisal code is identified by three major steps based on conversion of individual neuron’s spike trains into variability distribution of ISI, followed by its conversion to real-time self-information value and subsequent surprisal code at the cell-assembly level.

While traditional techniques used to decode stimulus identity or neuron’s tuning property (i.e. based on across-trial data-averaging methods, such as peri-event spike histograms) can be essential, such practice can also be intrinsically problematic because the brain is unlikely to use such trial-averaging methods to encode information in real-time (Figure 1B). As such, firing variability makes single neuron-based rate and temporal codes unsatisfactory in explaining how the brain actually achieves robust neural coding in real-time. This begs the questions of why and how neurons should perform real-time neural computation.

One of the most promising approaches in investigating neural coding is the study of cell assemblies. The concept of cell assembly – a group of neurons that fire transiently or sequentially – has been a cornerstone in brain research and often viewed as the computational primitives to encode an object, concept or memory engram. Yet, by and large, the identification of cell assemblies in both awake and sleep states has remained the state-of-the-art in systems neuroscience research (17, 22, 34-40). In part, this challenge is again due to enormous firing variability of individual neurons (41-44). From a signal-processing perspective, such variability (in either spike count or spike timing) would represent system noise, which has been shown to undermine reliable decoding of stimulus identities in real-time (42, 43). However, several interesting studies have also suggested that neuronal variability can be beneficial for boosting signal or serving as modulatory signals (45-50). More recently, large-scale recording studies have shown that neurons with similar tuning properties, termed neural cliques, can overcome individual variability by temporal summation and coordination of their joint activations (12, 20, 24, 40) (Figure 1C).

Here, we ask further the question of what type of design principles a neuron should use so that it can deal with such complex and variable synaptic input patterns while still using spikes to convey information in real-time and would be resistant to interferences. Our hypothesis regarding the role of neuronal variability in neural coding is based on the self-information theory. Specifically, we hypothesize that neuronal variability can be viewed as the self-information generator and expressor. Under this conceptual framework, neurons transmit their information by conforming to the basic logic of the statistical Self-Information Theory: spikes with higher-probability inter-spike-intervals (ISIs) contain less information, whereas spikes with lower-probability ISIs – termed variability-surprisal spikes – convey more information. In other words, real-time information is encoded not by changes in firing rates per se, but rather by variability-surprisal spikes (ISI patterns that have lower-occurring probability) (Figure 1D). These variability-surprisal spikes can be either positive-surprisals (when a neuron’s ISI becomes much shorter from typical, reflecting excitation) or negative-surprisals (when ISI becomes much longer than typical, reflecting inhibition). As such, these dynamic, transient surprisal-spikes can constitute real-time information packets to construct real-time temporally coordinated cell-assembly code.

The variability-surprisal based neural self-information theory makes several testable predictions: First, if neuronal variability acts as a self-information generator, this variability should remain similar across various brain regions. On the other hand, if neuronal variability reflects system noise, one would expect that variability would grow larger as information is transmitted from low subcortical structures to the high-cognition cortices. To differentiate these two scenarios, one can record large numbers of neurons from various cortical and sub-cortical regions in freely behaving animals. To facilitate systematic comparisons, one can initially focus on putatively classified principal cells after these units have been separated from fast-spiking putative interneurons and their variability distributions analyzed across these different regions. In addition, to minimize the potential state-dependent influence on neuronal variability, one can assess the spike datasets collected from the quiet awake state as animals rested in their home cage environments, as well as during various cognitive tasks.

One can characterize neuronal variability by using three well-defined statistics to describe quantitatively neuronal variability of a neuron’s ISI, - namely, a coefficient of variation (CV), skewness and kurtosis. In probability theory and statistics, CV is a standardized measure of dispersion of a probability distribution, and skewness is a measure of the asymmetry of a probability distribution, whereas kurtosis is a measure of the “tailedness” of a probability distribution. We would predict that principal cells in various brain regions should exhibit similar neuronal variability distributions. To further test the idea that neuronal variability serves as a self-information carrier, we would also predict that variability will diminish under the condition when both external and internal neural computation is artificially shut down (i.e. upon anesthesia). Pharmacological intervention experiments can be used to demonstrate that the shutting down of external and internal coding processes would indeed greatly reduced neuronal variability. If so, it would be consistent with the notion that spike variability reflects ongoing cognitive processing of both external and internal information.

One immediate and major implication of this “surprisal-spikes” concept is that it should enable researchers to identify a variety of cell assemblies. Overall, we suggest that this variability-surprisal-based, cell-assembly decoding (VCAD) strategy can consist of the following three major steps as follows (Figure 1D): The first step is to convert each neuron’s spike train into the probability distribution of ISI variability. One can be achieved by fit single neuron’s ISIs with a Gamma distribution model which can assign each neuron’s ISI with a probability. The second step is to convert the probability distribution of ISI variability into real-time self-information distribution for each ISI. These frequent ISI variations with high probability represent the low self-information or ground state (Figure 1D, the dotted gray curve in the mid-section of the Self-information plot) As a neuron increased its firing, it generates positive-surprisals (Figure 1D, the red curve inside the Self-information plot) as ISIs entered the left tail-zone of the distribution probability (a low-probability state). On the other hand, if the neuron’s firing is dramatically suppressed, negative-surprisals are generated (Figure 1D, the blue curve inside the Self-information plot) when ISIs shifted to the right tail-zone (also a low-probability state). Subsequently, a spike train emitted by a neuron can be transformed into surprisal-based ternary code (positive-surprisal, ground state, negative-surprisal) to describe the dynamic evolution in self-information states (Figure. 1D).

Subsequently, researchers should be able to uncover joint surprisal-spike patterns across simultaneously-recorded cells on a moment-to-moment basis. Blind source separation (BSS) methods, such as independent component analysis (ICA), can identify a set of independent information sources from simultaneously observed signals as structured patterns or relationships. Each independent signal source decoded by BSS would correspond to a distinct real-time activation pattern given by a cell assembly. To discover its functional meaning, one can compare each real-time activation temporal pattern with various other experimental parameters [such as the dynamic evolution of local field potential (LFP), the time points of stimulus presentations, videotapes of an animal’s behavioral state, actions, and corresponding locations, etc.]. Moreover, the top-ranking membership with the highest contribution weights in the cell assembly can be directly identified from demixing matrix W. This will allow researchers to assess quantitative membership information that other dimensionality-reduction-based, pattern-classification methods (i.e., principal component analysis or multiple discriminant analysis) could not provide. By further mapping cell-assembly activity patterns onto specific cell types and network states (51-53), we expect that researchers can gain greater insights into how neural code is generated within and across the evolutionarily conserved computational motifs (6-9, 14, 23, 54).

In short, as an illustration to the type of conceptual work listed by the Research Topic on Brain Activity Mapping^2.0^, we discuss a new hypothesis on how to crack the neural code. Specifically, we put forth a *neural self-information theory* that neuronal variability operates as the self-information generator and expressor to convey discrete quanta of information in the form of variability-surprisal spikes. Coordination of these surprisal spikes in space (across recorded cells) and time can be used to uncover various cell assemblies from various brain regions in an unbiased manner. The generality of this variability-surprisal spike concept can be demonstrated by identifying real-time cell assemblies processing internal states, external experiences - including continuous variables - and categorical variables. More importantly, this surprisal code can afford not only robust resilience to communication interference, but also biochemical coupling to energy metabolism, protein synthesis and gene expression at both synaptic sites and cell soma.

